# Paired Analyses of Nuclear Protein Targets and Genomic DNA by Single-Cell Western Blot and Single-Cell PCR

**DOI:** 10.1101/2025.03.29.646125

**Authors:** Ana E. Gomez Martinez, Trinh Lam, Amy E. Herr

## Abstract

Single-cell multimodal assays measure multiple layers of molecular information. Existing single-cell tools have limited capability to analyze nuclear proteins and genomic DNA from the same originating single cell. To address this gap, we designed and developed a microfluidic single-cell assay (SplitBlot), that pairs measurements of genomic DNA (PCR-based) and nucleo-cytoplasmic proteins (nuclear histone H3 and cytoplasmic beta-actin). To accomplish this paired multiomic measurement, we utilize microfluidic precision to fractionate protein molecules (both nuclear and cytoplasmic) from genomic DNA (nuclear). We create a fractionation axis that prepends a comet-like encapsulation of genomic DNA in an agarose molded microwell to a downstream single-cell western blot in polyacrylamide gel (PAG). For single-cell genomic DNA analysis, the agarose-encapsulated DNA is physically extracted from the microfluidic device for in-tube PCR, after release of genomic DNA from a molten agarose pallet (86% of pallets resulted in amplification of TurboGFP). For protein analysis, nucleo-cytoplasmic proteins are photocaptured to the PAG (via benzophenone) and probed in-situ (15 kDa histone H3 resolved from 42 kDa beta-actin with a separation resolution R_s_ = 0.77, CV = 76%). The SplitBlot reported the amplification of TurboGFP DNA and the separation of nuclear histone H3 and cytoplasmic beta-actin from the same single U251 cells engineered to express TurboGFP. Demonstrated here, Split-Blot offers the capacity for precision genomic DNA vs. protein fractionation for subsequent split workflow consisting of in-tube PCR and on-chip single-cell western blotting, thus providing a tool for pairing genotype to nuclear and cytoplasmic protein expression at the single-cell level.

**TOC Graphic:** 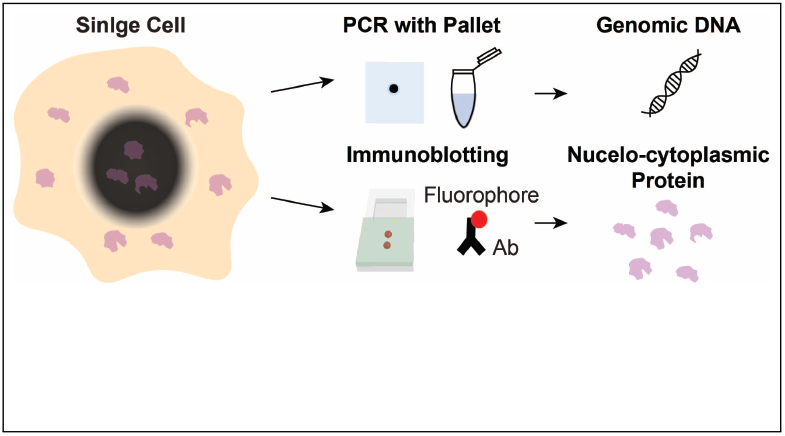

## Introduction

Advances in multimodal single cell assays directly link protein expression to the transcriptome, genotypes, or chromatin accessibility.^1,2^ As an example, CITE-seq^3^ and REAP-seq^4^ link assays of the transcriptome and cell-surface protein expression. In another example, DAb-seq detects surface proteins and sequences genomic DNA from the same single cells to track genotype-phenotype dynamics in leukemia tumors.^5^ Intracellular proteins and chromatin accessibility can be measured with ASAP-seq,^6^ while NEAT-seq^7^ adds a transcriptome mode to the intercellular protein and chromatin accessibility measurements. Paired same-cell, single-cell assay nucleic acid and protein assays elucidate gene regulatory networks and cell classification.^8,9^

Nuclear proteins – consisting of transcription factors, chromatin proteins, and nuclear structure proteins – are essential for regulating gene transcription and for signal transduction pathways.^10^ Nuclear proteins play crucial roles in cancer,^10^ neurodegenerative diseases, ^11^ cardiovascular diseases,^12^ and metabolic diseases.^13^ Single-cell intracellular protein expression can be analyzed by fixing and permeabilizing cells, followed by staining and analysis using flow cytometry, immunohistochemistry, or sequencing of oligo-labeled antibodies. However, oligo-labeled antibodies exhibit high background signals in the nucleus.^7^ In NEAT-seq, nuclear protein measurements are improved by blocking cellular components with ssDNA and blocking antibody oligo charges using E. coli ssDNA binding protein (EcoSSB). Another challenge in protein detection is that these methods require fixing and permeabilizing cells for the delivery of oligo-labeled antibodies. Proteins in fixed cells may experience antigen masking, likely due to epitope conformational changes or interactions within or between proteins.^14^ Antigen retrieval techniques can enhance protein detection,^15^ but they may also degrade epitopes.^14^ For nuclear protein analysis, fixation presents additional challenges. Challenges with fixation are exacerbated with nuclear protein targets because the highly concentrated nuclear compartment may limit antibody diffusion for probing.^16^ Further, chromatin crosslinking can mask epitopes.^17^ Both ASAP-seq and NEAT-seq perform nuclear protein measurements on permeabilized and fixed cells before immuno-probing with oligo-labeled antibodies, making salient the challenges of probing protein targets in fixed cells.

Microfluidic devices support high-throughput single-cell analyses. Droplet encapsulation of isolated single cells is suitable for ultra-high throughput analysis^2^ and is one of the most common approaches, whereas microwell isolation of single cells is well suited to categories of application where droplets struggle (i.e., scarce starting population of cells, large diameter cells). Both cell-isolation techniques are of interest, through, because reaction volumes are scaled down, yielding exceptional detection sensitivity and reduced cost per cell.

While multimodal assays with the ability to measure nuclear proteins (e.g. ASAP-seq and NEAT-seq) primarily employ droplet-based technologies, ^6,7^ microwell-based approaches are well suited to key applications, as mentioned. For example, from our own lab, the TriBlot performs paired analysis of cytoplasmic proteins and nucleic acids (mRNA or DNA) using microwells. The TriBlot has been optimized for paired analyses of ultra-low starting numbers of cells to study transitions in pre-implantation murine embryo development (i.e., blastomeres from two-cell and four-cell embryos) with protein isoform selectivity,^18–20^ an application that is inaccessible to highthroughput droplet-based approaches. Despite these advancements, a larger variety of tools designed to seamlessly analyze nuclear proteins and DNA from the same single cell would be welcomed.

In this study, we pair same-cell measurements of nuclear protein expression with a DNA measurement. Indexed to the same originating cell, microwell-based cell isolation allows joint analyses by single-cell western blot of nuclear proteins and single-nucleus PCR. To apply the western blot and PCR to the same starting cell, we designed a composite agarose-PAG device that facilitates in-microwell whole-cell lysis with electrophoresis into the agarose-PAG composite separation lane. Agarose abuts each microwell and physically retains DNA during electrophoresis, while nucleo-cytoplasmic proteins electrophoreses through the agarose and into the PAG protein-separation lane (Split-Blot). Then each pallet laden with DNA (in the microwell) is mechanically released from the planar device, allowing for independent PCR-based DNA analysis from the free pallet and immunoprobing in the PAG region, both results indexed to the same single cell. Split-Blot simultaneously analyzes DNA and nucleo-cytoplasmic protein from the same origination cell in an array, as evidenced by the amplification of TurboGFP DNA and size-based separation of beta-actin and histone H3.

## Results and Discussion

### Design of a Multimodal Single-Cell Assay

In order to make paired multiomics measurements of DNA and proteins from the same single cell possible, we designed and developed a single-cell assay to perform (i) precision singlecell fractionation of nuclear DNA from nucleocytoplasmic proteins (total proteins) and (ii) a matched workstream analysis of single-cell DNA with PCR and protein with single-cell western blotting. For brevity, we refer to this paired single-cell PCR and western blot assay as SplitBlot. To demonstrate the paired analyses, we designed a planar microfluidic device that accomplishes precision fractionation of the nuclear DNA compartment from the protein targets spanning the nucleo-cytoplasmic compartments using a series of stacked molecular-mass specific sieving materials and an applied electric field. In essence, we prepend a comet assay (large pore-size agarose molecular-sieving matrix)^21–23^ onto a western blot (small pore-size PAG molecular-sieving matrix)^24,25^ after wholecell lysis, and then physically separate the two matrix regions after single-cell capture of DNA and protein in each region, respectively.

This SplitBlot fractionation axis consists of a DNA-immobilizing comet region at the head of a protein-immobilizing western blot region (Figure 1a). The large-to-small pore-size pattern results in DNA stacking in the microwell-abutting agarose region while smaller molecules (i.e., proteins) electromigrate through the agarose region and into the PAG region for SDS polyacrylamide gel electrophoresis (PAGE), where protein targets are resolved by differences in molecular mass. Agarose retains the long DNA molecules,^21–23^ thus forming the basis for the immobilization and subsequent physical transfer of DNA to off-chip analysis; whereas, after PAGE separation, protein targets are retained in the PAGE region by photo-capture to benzophenone in the PAG. PAG pore diameters are ∼ 10 nm^26^ and agarose pore diameters are ∼ 100 nm.^27^ Each glass device houses a 5 × 10 array of the SplitBlot devices.

**Figure 1:**
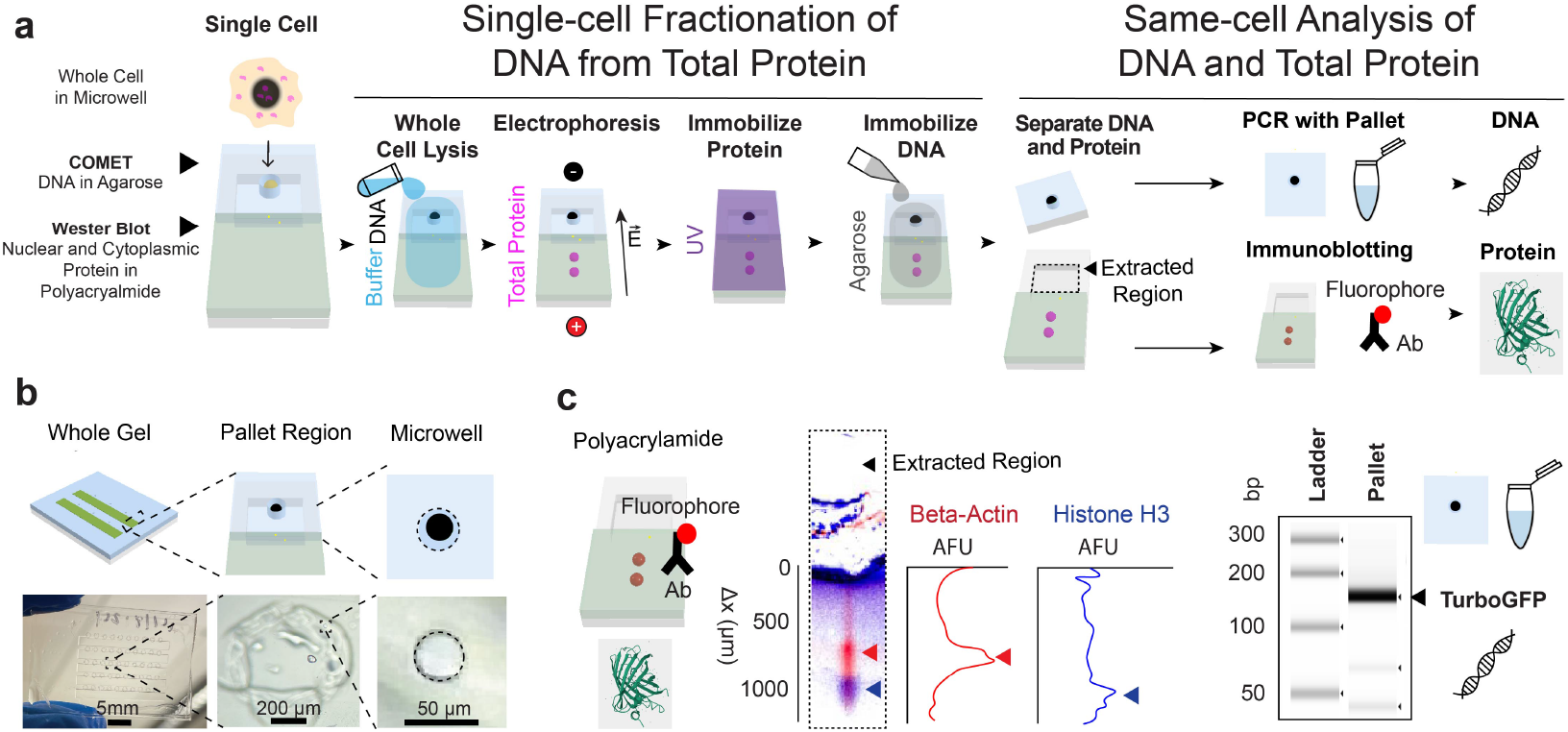
The SplitBlot assay uses a composite hydrogel and DNA extraction to perform paired analysis of DNA and intracellular proteins from the same originating cell. (a) Schematic of the workflow for SplitBlot shows that single-cell DNA and total proteins (both cytoplasmic and nuclear) are separated to perform multimodal measurements. The composite hydrogel consists of agarose regions containing a microwell for DNA capture and a PAG region for protein capture. Whole cells are lysed, and electrophoresis separates DNA and proteins in the composite hydrogel, where each molecule is immobilized. Then agarose pallets are extracted to perform PCR and the PAG is immunoprobed. (b) Schematic and brightfield photographs of the composite gel architecture illustrate that the composite gel has multiple lanes, each lane has a pallet region outlined by an engraved glass region, and each engraved glass region contains an agarose microwell. (c) Micrographs of immunoblotted PAG and of the agarose gel electrophoresis of the PCR product indicate that the expression of nuclear protein histone H3 (blue) and cytoplasmic protein beta-actin (red) can be measured while performing PCR of TurboGFP from the same single cell. Antibodies preferentially partition into edges of the PAG lanes, resulting in higher background in this region.

The SplitBlot comprises 6 sequential steps (Figure 1a). First, single cells from a cell suspension are sedimented into microwells located in each agarose region. Second, once isolated in a microwell, each isolated whole cell is chemically lysed using a dual lysis and electrophoresis buffer. Third, as cell lysis completes, electrophoretic-fractionation of the nuclear DNA fraction vs. nucleo-cytoplasmic protein fractions is initiated through application of an electric field physically aligned with the fractionation axis (i.e., microwell-in-agarose region for DNA capture, followed by PAG region for protein PAGE and immobilization). The protein electromigrates through the large-pore-size agarose comet region and into the smaller-poresize PAG region, while large molecular mass DNA remains immobilized in the agarose region. Fourth, once in the PAG region, proteins resolve based on differences in molecular mass (protein sizing) and are blotted (immobilized) through brief UV-light activation of benzophenone, which was incorporated into the PAG during gel polymerization. Fifth, DNA is sealed into the comet region through application of an agarose lid, which immobilizes DNA due to the negligible diffusivity of DNA in agarose. Sixth, each agarose comet region is mechanically released from the planar device as a DNA-laden agarose gel pallet. Release of the pallet is accomplished using a razor cut along an engraving line with subsequent mechanical release. After release, each agarose pallet is placed in a PCR tube and melted to liberate DNA into solution, allowing single-cell PCR to proceed. This completes the DNA analysis step of the multiomics SplitBlot assay. Concurrently, the PAG region still affixed to the planar glass substrate is immunoprobed to report presence and expression level of proteins from the cytoplasmic and nuclear compartments. Thus, completing the protein analysis step of the SplitBlot assay. Both the PCR and protein measurements are linked to the originating cell owing to spatial layout of the array, where spatial indexing of each gel region corresponds to an originating fractionation axis location and, thus, originating microwell.

The fractionation axis architecture was designed as composite agarose-PAG regions to allow for two concurrent but distinct analysis streams for each single cell (Figure 1b), namely (1) the extraction of each DNA-laden agarose region as a pallet for off-chip PCR analysis and (2) retention of the PAG region on the planar glass device to support intracellular protein analysis by single-cell western blot. As described in detail in subsequent sections, the SplitBlot assay reports protein expression — including, importantly, expression of nuclear proteins such as histone H3 — while performing same-cell PCR of TurboGFP, as a demonstration of nuclear-compartment analysis of protein and DNA from the same originating cell but via different analytical modalities (Figure 1c).

### Single-Cell DNA Analysis from a Detachable Agarose Gel Pallet

Inspired by classical and microarray-based comet assays,^21,23^ we hypothesized that the molecular mass of genomic DNA would prohibit substantial diffusive loss of the DNA from each microwell both (i) during the whole-cell lysis and electrophoresis periods and (ii) during subsequent pallet extraction and transfer periods owing to the application of an agarose lid to encase the liberated genomic DNA in a pallet-like feature. The encapsulating lid protects the DNA from loss by both convection (during washing and handling) and diffusion. To create an estimate of characteristic diffusion times for molecular loss, we employed the reptation model for DNA diffusion in a hydrogel^28^ and the Stokes-Einstein relation for diffusion in free solution. The genomic DNA is in free solution during lysis and electrophoresis. Based on the diffusion coefficient of DNA in free solution, 50 kb dsDNA should diffuse 42 *µ*m (microwell height) in 21 min, which is slower than the lysis and electrophoresis timescale of ∼1 min. Subsequently, application of a 58-*µ*m thick agarose lid (total gel height of 100 *µ*m) would see the 50-kb dsDNA diffuse out of 1% agarose in ∼80 days. Thus, we concluded that the agarose encapsulation would be effective at immobilizing DNA for the downstream handling steps.

In our design of the genomic DNA analysis, we sought to devise an approach to toggle between two discrete functions: (i) encapsulation and immobilization of DNA for transfer from the microfluidic device to the tubes and (ii) release of the DNA for the PCR analysis step. To achieve this toggling of function, we explored designs using the temperature-sensitivity of normal melting temperature agarose, with a temperature rise triggering the transition of a solid agarose pallet immobilizing DNA (temperature point: 36 °C) to a molten agarose state (temperature point: 90 °C) suitable for release of DNA into a surrounding reservoir of free solution (PCR tube).

To understand the utility of agarose formulations for the first function, we compared the immobilization of DNA in normal melting temperature agarose with and without a low melting temperature agarose lid. Single U251 cells were settled into microwells, excess cells are washed off the open-fluidic device, leaving a distribution of microwell occupancies (25% of agarose microwells with single-cell occupancy; 9% with multiple cell occupancy; 66% of microwells empty in Figure 2a). This is comparable to the distribution of microwell occupancies in a PAG (19% of microwells with single-cell occupancy; 3% with multiple cell occupancy; 78% of microwells empty). After single-cell settling, cells were chemically lysed in situ (20 s), and electrophoresis was performed (40 s, 40 V/cm). Molten low melting temperature agarose at 37 °C was placed on top of the composite hydrogel and gelled at room temperature. DNA was stained with 1x SYBR Green I to determine the location of DNA. We observed that after electrophoresis for 40 s, DNA partially injected into the solid agarose region. With an agarose lid, DNA was immobilized in the microwell, which is suitable for the design goals of agarose in the SplitBlot assay (Figure 2b). Without the agarose lid, DNA diffused from the microwell. This analysis suggests that intact genomic DNA is too large to inject completely during the electrophoresis period. Based on this analysis, an agarose encapsulation lid was incorporated into the SplitBlot assay to retain DNA in the microwells for release and pallet transfer for off-chip PCR.

**Figure 2:**
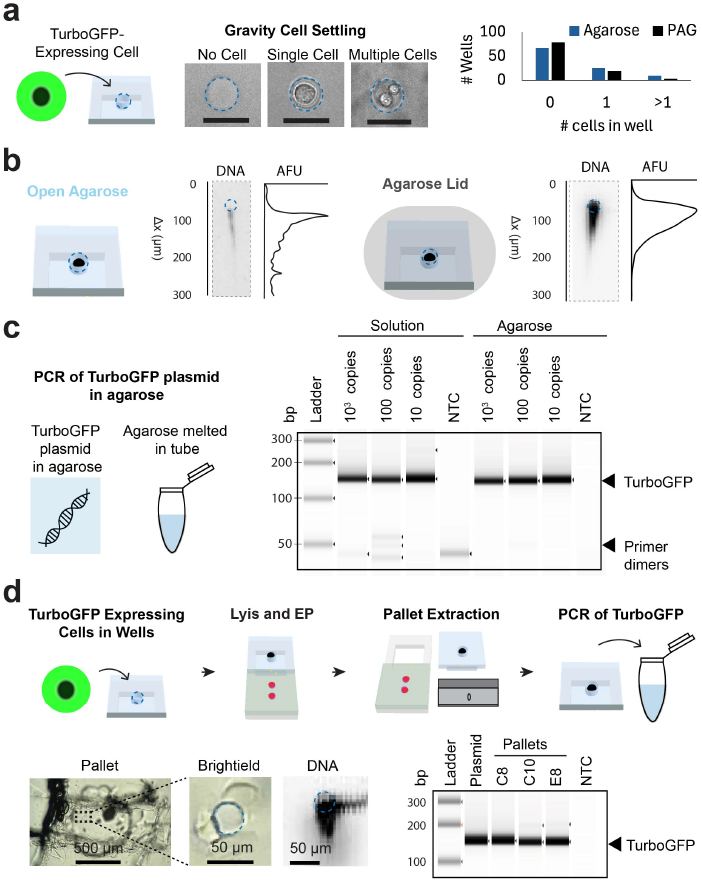
Single-cell DNA analysis uses agarose to contain DNA within the pallets for downstream transfer, release, and PCR. (a) Brightfield micrographs of agarose microwells and the bar graph of microwell occupancy demonstrate that single U251 cells were gravity settled into agarose microwells (n = 100 microwells). Scale bar = 50 *µ*m. (b) Micrographs and electropherograms of DNA in agarose illustrate that a low melting temperature agarose lid immobilized genomic DNA that was not injected in the PAG during 40 s electrophoresis. (c) Micrographs of the agarose gel electrophoresis of the PCR product confirm the release of a serial dilution of DNA, specifically TurboGFP plasmid, from agarose. TurboGFP was amplified both in solution and from plasmids released from agarose at the lowest tested concentration of 10 copies/*µ*L. (d) The schematic and brightfield photographs of the gel convey the workflow for performing PCR from single U251 cells engineered to express TurboGFP. Cells were isolated in microwells, and agarose pallets (3.5% NMTA) were mechanically released from glass with a razor blade and placed in PCR tubes. Micrograph of DNA in microwell confirmed isolation of DNA and micrographs of the agarose gel electrophoresis of the PCR product show TurboGFP amplified from agarose pallets.

To understand the utility of agarose formulations for the second function, we examined the release of DNA from the normal melting temperature agarose hydrogel. We did this by comparing the amplification of a serial dilution of a TurboGFP plasmid embedded in agarose to the amplification of a serial dilution of TurboGFP in solution (Figure 2c). Plasmids were released from normal melting temperature agarose by melting the pallet at 100 °C and diluting the agarose in water to prevent re-gelling. We observed that the lower limit of detection for the amplification of TurboGFP (10 TurboGFP plasmid copies per pallet) was the same for the in-solution DNA control and for DNA release from agarose by temperature-toggling the agarose to a molten state. This supports the design assertion that the agarose pallet can release DNA from normal melting temperature agarose for PCR when heated.

Next, we verified the ability to perform PCR from mechanically released agarose pallets consisting of a normal melting temperature base with a low melting temperature agarose lid. Single cells engineered to express TurboGFP were settled in 3.5% normal melting temperature agarose microwells, whole-cell lysis and electrophoresis was performed, 3.5% low melting temperature agarose lid was gelled, and DNA was stained to identify pallets with DNA in microwells. The agarose pallets were released by sectioning the agarose pallet along the engraving on the glass using a razor (Figure 2d). Then, the agarose pallets were sheared off the glass and transferred into PCR tubes, one pallet per tube. We observed TurboGFP bands in the micrographs of the agarose gel electrophoresis of the PCR product, signifying that we amplified TurboGFP from agarose pallets containing single-cell DNA (n = 3 single-cell pallets). The observations verified that this extraction method allowed for transfer of DNA and retained quality DNA for PCR.

### Single-Cell Immunoblotting from PAG-Fractionated Nuclear and Cytoplasmic Proteins

To ascertain the SplitBlot assay’s capability to detect nucleo-cytoplasmic protein targets — while subjecting genomic DNA to PCR from that same cell — we next examined the electromigration of TurboGFP protein in the agarose-PAG fractionation axis. U251 cells engineered to express TurboGFP were utilized as a case study. We used time-lapsed fluorescence imaging and resultant micrographs to determine the TurboGFP location during lysis and electrophoresis. We observed that TurboGFP is retained in the microwell during 20 s of whole-cell chemical lysis. Upon application of an electric field (40 V/cm), we observed that TurboGFP migrates with a velocity of 30.4 *µ*m/s along the length of the composite gel fractionation axis (Figure 3a). We confirmed that, during lysis, protein remained in the microwells molded in agarose, and, during electrophoresis, proteins electromigrated through the comet agarose region and into the PAG region, transitioning from a molecular fractionation mechanism (genomic DNA vs. protein) and into a protein sieving mechanism yielding a size-based protein separation in the PAG region.

**Figure 3:**
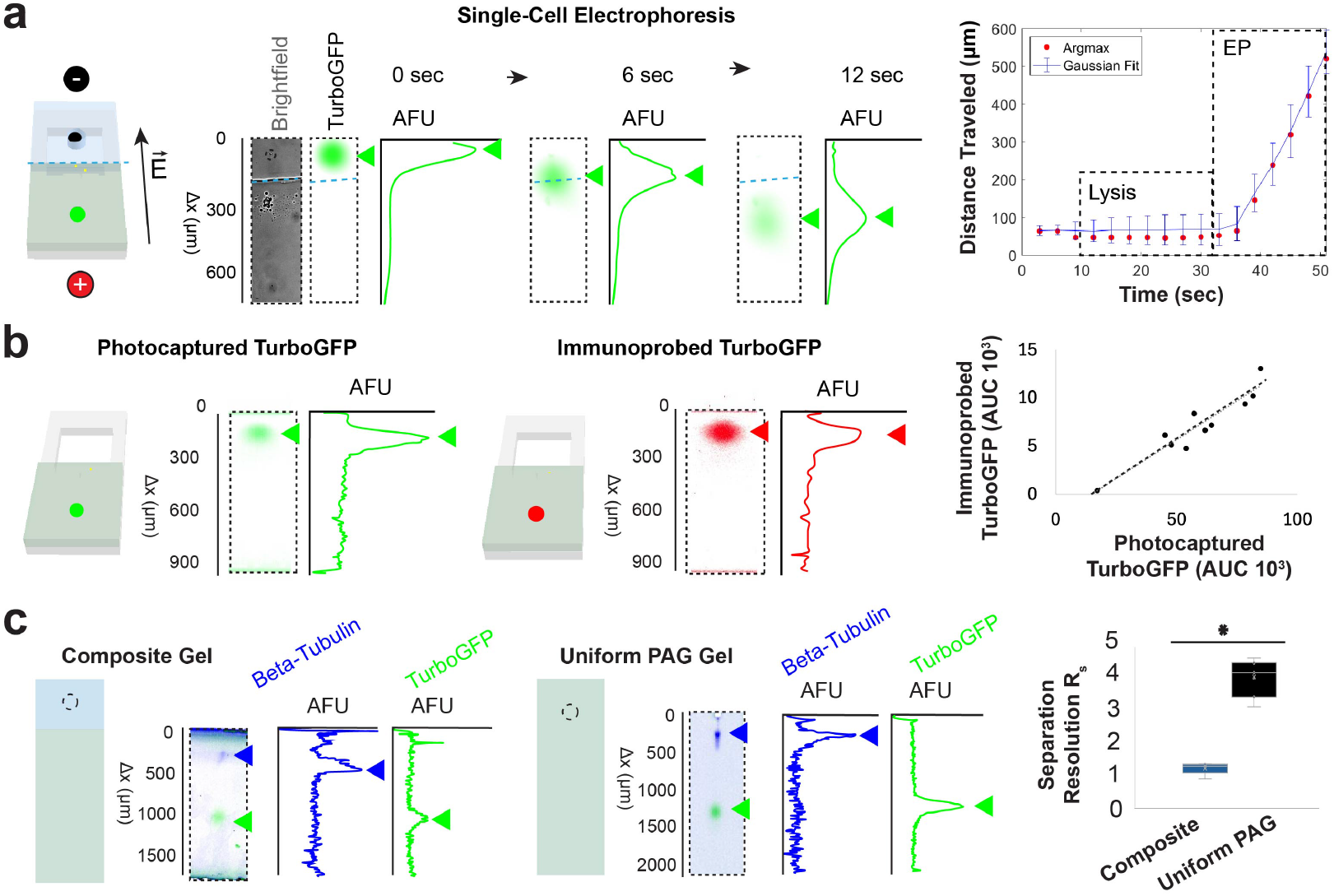
The composite gel enables protein migration, protein separation, and immunoprobing of photocaptured proteins. (a) Electropherograms of TurboGFP and plot of TurboGFP location during lysis of single U251 cells engineered to express TGFP and electrophoresis of TurboGFP in a composite gel demonstrate that TurboGFP migrated in the composite gel with an average speed of 30.4 *µ*m/s during electrophoresis at 40 V/cm. The Gaussian fit for the plot of distance over time shows the location and the band width of TurboGFP while Argmax is the location of the maximum intensity. (b) A scatterplot of the AUC for photocaptured TurboGFP and immunoprobed verified that there is a correlation between the AUC of photocaptured and immunoprobed protein in the composite gel. (c) A boxplot of separation resolution for TurboGFP (27 kDa) and beta-tubulin (55 kDa) in a composite and uniform PAG indicate proteins were separated in both the composite and uniform PAG.

After successful protein sizing in the PAG region, we evaluated immunoprobing in the PAG region of the SplitBlot device by determining the relationship between immunoprobed (i.e., detected by fluorescently labeled primary antibody) TurboGFP protein and natively fluorescent photo-captured TurboGFP protein (Figure 3b). Immunoprobed TurboGFP protein and photocaptured TurboGFP protein showed a positive correlation (Pearson Correlation Coefficient 0.94, p-value < 0.05, n = 10 cells), thus verifying the capability of the SplitBlot assay to perform immunoprobing in the PAG region of the composite hydrogel.

Lastly, to establish the analytical capability of the single-cell western blot workstream of the SplitBlot assay, we scrutinized the separation capabilities of the composite gel (1% agarose and 8%T (29:1) PAG) compared to an 8%T (29:1) uniform PAG (Figure 3c). Here, cells engineered to express TurboGFP were isolated in microwells, we performed lysis and PAGE, and beta-tubulin was immunoprobed. The separation resolution (R_s_) after 50 s electrophoresis at 40 V/cm for TurboGFP (27 kDa) and betatubulin (55 kDa) was 1.2 (CV = 16%; n = 5 cells) in a composite hydrogel and 3.0 (CV = 19%; n = 33 cells) in a uniform PAG. Based on the location of TurboGFP and beta-tubulin, the range of molecular masses expected to be photocaptured in this composite gel is 12 kDa-79 kDa. In the composite gel, the average peak width of TurboGFP was 250 *µ*m (CV = 47%; n = 5 cells), and average peak width of betatubulin is 70 *µ*m (CV = 21%; n = 5 cells). In a uniform PAG, the average peak width of TurboGFP is 55 *µ*m (CV = 4.9 %; n = 33 cells) and the average peak width of beta-tubulin is 165 *µ*m (CV = 24%; n = 33 cells). Peak dispersion in the gel is due to injection and diffusion. A smaller molecular mass protein TurboGFP diffuses in agarose, causing band broadening that is not rescued by stacking, resulting in a 4.5X greater peak width in a composite gel compared to in a uniform PAG. Dispersion even during stacking is evidenced by analyzing the ratio of peak width for TurboGFP after crossing the agarose-PAG interface to the peak width before crossing the interface (ratio = 1.57). However, beta-tubulin, a larger molecular mass protein, experiences less diffusive losses and stacks in the composite gel, resulting in a 2.4X smaller peak width in a composite gel compared to in a uniform PAG. Taken together, separation resolution is lower for this set of proteins in composite gels than in PAG due to the dispersion of lower molecular mass protein in the agarose gel. Nevertheless, TurboGFP and beta-tubulin were resolved in both the composite and uniform PAG hydrogel.

### Paired Genomic DNA and Nucleo-Cytoplasmic Protein Measurements

We next examined the performance of the Split-Blot assay in implementing paired analyses of genomic DNA and nuclear proteins from the same starting U251 cells. We again utilized U251 cells engineered to express TurboGFP, and gravity settled individual cells into microwells formed in 3.5% normal melting temperature agarose. To fractionate the genomic DNA vs. protein, we performed whole-cell lysis for 20 s and PAGE at (70-80 s at 40 V/cm) then encapsulated the entire gel in 3.5% low melting temperature agarose. To verify the immobilization of genomic DNA, the DNA was stained and pallets with DNA in microwells were selected for physical extraction (Figure 4a). To demonstrate detection of genomic DNA, we performed pallet transfer and DNA release as described above, followed by in-tube PCR. We observed TurboGFP bands in the micrographs of agarose gel electrophoresis of PCR products (Figure 4b). To demonstrate detection of proteins – including nuclear proteins – from the same cell – beta-actin (42 kDa) and nuclear histone H3 (15 kDa) were subjected to single-cell protein fractionation and western blotting. We found beta-actin and histone H3 were resolved at R_s_ = 0.77 (CV = 76%; n = 7 lanes) in the PAG region (Figure 4c). Taken together, the SplitBlot assay was capable of amplifying TurboGFP from genomic DNA and reporting by western blot the presence and expression of the cytoplasmic protein beta-actin and the nuclear protein histone H3 from the same originating cell.

**Figure 4:**
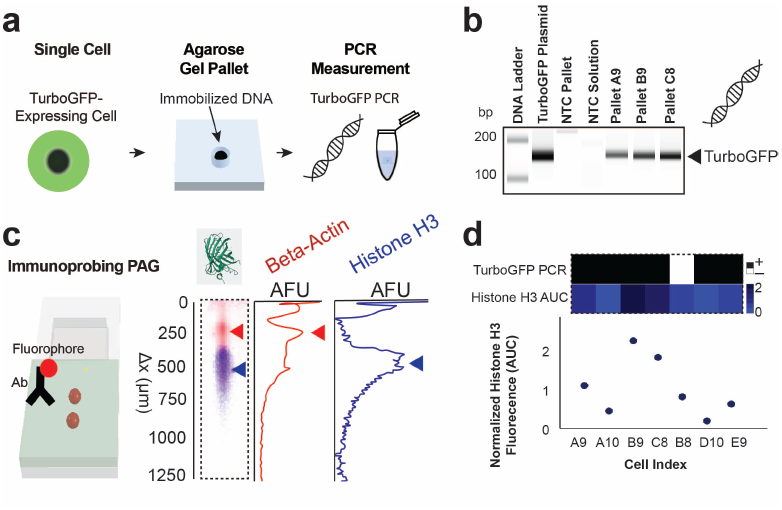
The SplitBlot detects nuclear proteins and cytoplasmic proteins while performing PCR from the same single cell. (a) Schematic illustrates key steps in the workflow to perform PCR from single cells. We performed singlecell PCR by isolating single cells in agarose microwells and extracting the agarose pallet laden with DNA. (b) Micrographs of agarose gel electrophoresis of PCR products evidence the presence of the TurboGFP amplicon from single cells. (c) Micrographs of immunoblotted PAG show beta-actin (42 kDa) and histone H3 (15 kDa) protein bands were separated. (d) Heatmap of PCR results and normalized histone H3 expression demonstrate that we can detect nuclear histone H3 and cytoplasmic betaactin in PAG while amplifying TurboGFP from the same single cells. Intracellular protein expression is linked to PCR results based on the pallet and lane indexed in the composite gel.

Using the single-cell western blot workstream, we analyzed the histone H3 expression in each of these single cells. We estimated histone H3 protein expression by normalizing histone H3 area-under-the-curve (AUC) to beta-actin AUC. Histone expression was expected to depend on the cell cycle, with expression peaking during S-phase and correlating with genome content.^29^ The mean normalized histone expression was 1.0 with a CV of 62% (n = 7 cells). As expected, we observed a distribution of histone H3 expression levels across the cells analyzed.

Lastly, we evaluated the efficiency of the DNA extraction and transfer method. We analyzed the PCR results of pallets linked to protein bands based on the array index (Figure 4d). We found that six out of seven pallets linked to a protein measurement resulted in amplification of TurboGFP from genomic DNA, suggesting that the SplitBlot assay reported genomic DNA (from the nucleus) and protein targets from both the nucleus and cytoplasm in 86% of cells assayed by western blot.

## Experimental

### Chemicals and Materials

Solution of 30% (w/v) (29:1) acrylamide/bis acrylamide (A3574) N, N, N’, N’tetramethylethylenediamine (TEMED, T9281), ammonium persulfate (APS, A3678), 3-(Trimethoxysilyl) propyl methacrylate (98%, 440159) were purchased from Sigma-Aldrich. N-[3-[(3-Benzoylphenyl)-formamido]propyl] methacrylamide (BPMAC) was synthesized by PharmAgra Laboratories. Dual lysis/electrophoresis buffer is composed of X0.5 Tris-glycine from Bio-Rad (1610734), 0.5% sodium dodecyl sulfate (SDS) from SigmaAldrich (L3771), 0.25% sodium deoxycholate from Sigma-Aldrich (D6750), 0.1% Triton™ X-100 from Sigma-Aldrich (X100-100ML). Harsh stripping buffer is composed of 62.5 mM Tris-HCl (Teknova, T1568), SDS at 2% (wt/vol), and 0.8% *β*-mercaptoethanol (Sigma-Aldrich, M3148). Gel Slick^®^ solution (50640) was purchased from Lonza. 10x Tris/Borate/EDTA buffers (TBE, AM9863), normal melting temperature agarose (NMTA, 16500500), low melting temperature agarose (LMTA, 16520100), 10 000x SYBR Green I (S7563), UltraPure™ DNase/RNase-Free Distilled Water (10977015) were purchased from Invitrogen. Tris Buffered Saline with Tween^®^ 20 (TBST 10X) was purchased from Cell Signaling Technology (9997S). Allyl agarose (AA8003) was purchased from Lucidant Polymers. Phosphate-buffered saline (PBS, pH 7.4, 1001002) was purchased from Thermo Fisher Scientific. Taq PCR Kit (E5000S) was purchased from New England Biolabs. DNA Clean and Concentrator-5 (D4013) were purchased from Zymo Research. Qubit™ dsDNA HS Assay Kit (Q32854) was purchased from Thermo Scientific. Agilent Technologies high sensitivity D1000 reagents (UFUCOP-5067-5585) and high sensitivity D1000 screentape (UFUCOP-5067-5584) were purchased form Neta Scientific. TurboGFP primers were purchased from Integrated DNA Technologies.

### Cell Culture

U251 cells were stably transduced with TurboGFP via lentiviral infection (multiplicity of infection 10). U251 cells were cultured in DMEM, high glucose, GlutaMAX™ Supplement (10566-016) with 1% penicillin/streptomycin (15140122, Life Technologies), BenchMark™ FBS (100-106, Gemini Bio-Products), 1× MEM nonessential amino acids (11140050, Life Technologies), and 1 mM sodium pyruvate (11360-070) in an incubator at 37 °C with humidified 5% CO_2_ air.

### Device Fabrication

SU-8 wafers were fabricated using photolithography as previously reported.^24^ The SU-8 wafer with posts was used in a double molding process to obtain PDMS microwells and then PDMS microposts (SI Figure 1). Each micropost used to fabricate the array of agarose microwells was 32 *µ*m in diameter, 42 *µ*m high, the pitch was 1.9 mm (transverse) by 3 mm (axial). Another SU-8 wafer was fabricated to mold PAG lanes. The SU-8 mold to fabricate PAG regions was 42 *µ*m high, the separation lanes were 2 mm in length axially with 1 mm spacing between lanes. Previously, we introduced a multimodal assay (called TriBlot)^20^ that contained 2.40 microwells/cm^2^ and, here, SplitBlot contains 14.6 microwells/cm^2^, a factor of 6 increase in the number of microwells per area. Glass slides treated with 3-(trimethoxysilyl)propyl methacrylate as previously reported.^24^ Then, glass slides were coated in molten 1% allyl agarose dissolved in water by spreading 1 mL of the molten agarose using the pippet tip, removing excess agarose by tilting the glass 45 degrees and wicking excess agarose into kimwipe. The agarose was air dried for >2.5 h. The glass slide was laser engraved to create markings that subsequently guide perforation and release of each pallet, containing a microwell in agarose, by razor cutting. Pallet regions of 1 mm in length were laser engraved on the agarose coated side of the glass (SI Figure 2). The CO_2_ laser (Full Spectrum Laser with a maximum power of 40 W, MUSE-40) engraving setting were 100% speed, 1 pass, and 2% power. The SU-8 wafer for PAG molding was coated with 1 mL of Gel Slick^®^ solution, wiped clean with a Kimwipe^®^, and dried with a nitrogen stream. PAG precursors with 3 mM BPMAC, 0.08% TEMED, and 0.08% APS, 1x Tris-Glycine were degassed 10 min, were molded, and were polymerized for 15 min in a humidity chamber (water vapor is created due to the evaporation of water from a water-soaked KimWipe). An array of 32-*µ*m diameter microwells were molded into agarose (with a PDMS mold) on top of the laser engraved glass region. The PDMS was coated in Gel Slick^®^ solution by adding 1 mL of Gel Slick^®^, removing excess Gel Slick^®^ by tilting, using nitrogen stream to move Gel Slick^®^ droplets on the PDMS surface in circles until droplets disappear. The PDMS mold and PAG bonded to engraved glass were each heated to 60 °C, molted normal melting temperature agarose was added to the mold, and the pallet regions on the glass were aligned to the microposts of the PDMS. Agarose was kept molten during alignment by placing a hotplate at 50 °C under a stereoscope. Fiducial markers on each layer guided alignment (see.dwg file in SI). The glass slide was taped down onto the assembly and a weight (∼ 20 g) was applied to the top to create a thin agarose layer. Agarose was gelled at 4 °C for 15 min and dehydrated 5 min before demolding. The composition of the PAG (for the separation of beta-actin and histone H3) and the agarose pallet (for encapsulation and extraction of DNA) was 7%T (29:1) PAG and 3.5% agarose, respectively. The composition of all other composite gels was 1% agarose and 8%T (29:1) PAG and the uniform PAG was 8%T (29:1). A guide to troubleshooting the fabrication and assay is in SI Table 1.

### Single-Cell Western Blot

U251 cells engineered to express TurboGFP were gravity settled in normal melting temperature agarose microwells by prewetting agarose with 200 *µ*L of PBS and pipetting 200 - 500 *µ*L of strained cells at concentration of 1×10^6^ cells/mL. Cells gravity settled for 15 min and unsettled cells were washed away by placing gel at 45 degrees and flowing 1 mL of PBS on each side of the gel. The composite gel was placed in an electrophoresis chamber with 100 *µ*m thick tape on edges, and the ice-cold dual lysis/electrophoresis buffer was carefully poured onto the chamber. Dual lysis/electrophoresis buffer is composed of 0.5x Tris-glycine, 0.5% SDS, 0.25% sodium deoxycholate, 0.1% Triton™ X-100. ^24^ We performed whole-cell lysis for 20 s and electrophoresis for 40-80 s at 40 V/cm. Photocapture of protein was accomplished with a UV light source (100% power, 45 s, Lightningcure LC5, Hamamatsu).^24^ For the SplitBlot assay, the excess buffer was removed with pipette, and the composite gel was coated in thin low melting temperature agarose lid by adding 300 - 500 *µ*L molten 3.5% low melting temperature agarose in PBS at 37 °C and adding a Gel Slick^®^ solution treated glass slide on top, pressing down slightly so the glass contacted the 100 *µ*m thick tape rails. The agarose was gelled for 10 min at room temperature, then rehydrated with PBS and stored in PBS until pallets were stained and extracted.

### Pallet Extraction and Single-Cell PCR

DNA was stained with 1x SYBR Green I for 10 min and imaged while hydrated to determine which pallets to extract based on the visible presence of stained single-cell DNA, meaning a single comet in the pallet region. The composite gel was rinsed in deionized water. Agarose was maintained in a hydrated state before removing all pallets because dehydrated pallets cannot be sheared off glass. However, excess water can lead to the loss of pallets in the water layer. Therefore, the agarose was kept semi-hydrated by adding ∼ 100 *µ*L deionized water every ∼ 5 min. Using brightfield microscopy, the perimeter of the pallet was cut with a razor guided by the glass engraving as a template. The razor was used to carefully shear the agarose pallet off the glass surface and one pallet was placed in each 50 *µ*L volume PCR tube with 38.5 *µ*L distilled molecular-grade water using tweezers. The tube was labeled with the corresponding index of the array position. Out of 22 pallets extracted, 100% of these could be moved into the PCR tube. The individual pallets were melted at 100 °C by mixing the solution with a pipette every 3 min for 20 s and repeating until the pallet melted (<10 min). Tubes containing melted pallets were stored at-20 °C. Positive controls were 100 copies of TurboGFP plasmids. We added 11.5 *µ*L of master mix to each tube. The final concentration of PCR mixture was 500 nM each primer (5’GACAGCGTGATCTTCACCGA and 5’ TCCACCACGGAGCTGTAGTA), 1x Standard Taq Reaction Buffer, 200 nM dNTP, 25 units/mL Taq polymerase. The first stage of thermal cycling was 95 °C for 4 min, the second stage (denature at 95 °C for 30 s, annealing at 55 °C for 30 s, extension at 68 °C for 45 s) was 45 amplification cycles, and a final stage was 72 °C for 5 min. The heated lid was set to 105 °C. The PCR products were concentrated with a DNA Clean and Concentrator-5 kit into 6.5 *µ*L. A Qubit 4 Fluorometer was used to measure the concentration of the PCR product and PCR product was diluted in water to 2 ng/*µ*L in order to measure the PCR product sizes using the 4150 TapeStation System with the high sensitivity D1000 reagent and screentape.

### Single-Cell Immunoprobing

After pallets were extracted, the composite gel was washed in 1x TBST for 30 min, then rinsed in deionized water. Excess agarose was delaminated with tweezers. Dehydrated agarose cannot be removed but excess water on the gel does not allow for distinction of agarose vs PAG by visual inspection so the gel was semi-hydrated during the removal of agarose by adding ∼100 *µ*L deionized water every ∼5 min. Immunoprobing protocol was as previously described^24^ by placing the gel on top of a glass plate with rails and the antibody solution, and then moving the gel of the gel off the rails and removing the rail, one rail at a time. Previously, two rails were used but, here, a single rail was used to avoid damaging the gel on a second rail. The antibody cocktail volume was 60 *µ*L per half slide. We incubated the primary antibody for 2 h, washed for 1 h in 1x TBST (buffer exchanged every 30 min), incubated the secondary antibody for 1 h, and washed 1 h in 1x TBST (buffer exchanged every 30 min). The primary antibody cocktail for the multimodal measurements was rhodamine labeled anti-beta-actin (1:5 dilution, 12004163) and rabbit anti-histone H3 (1:10 dilution, ab1791) in 2% BSA in 1x TBST. The secondary antibody was donkey Alexa Fluor Plus 647-conjugated anti-rabbit (1:20 dilution, A32795) in 2% BSA in 1x TBST. The primary antibody cocktail for analyzing separation resolution was rabbit anti-beta-tubulin (1:10 dilution, ab6046) in 2%BSA in 1x TBST and the secondary antibody was donkey Alexa Fluor Plus 647 conjugated anti-rabbit (1:20 dilution, A32795) in 2% BSA in 1x TBST. TurboGFP was probed with DyLight 594 labeled mouse anti-TurboGFP (1:5 dilution, TA150024) in 2% BSA in 1x TBST. Antibodies are stripped from the PAG with a 30 min incubation at 55 °C in a harsh stripping buffer. Harsh stripping buffer is composed of 62.5 mM Tris-HCl, SDS at 2% (wt/vol), and 0.8% *β*-mercaptoethanol. The PAG was re-probed after a 1 h wash in 1x TBST.

### Imaging

Fluorescence from each protein band was imaged with 5 *µ*m/pixel spatial resolution using a GenePix 4300/4400 Microarray Scanner from Molecular Devices (San Jose, CA). Cells in microwells and time-lapse micrographs of TurboGFP were imaged with a 10× magnification objective (Olympus UPlanFLN, NA 0.45, Tokyo, Japan). The Olympus IX71 inverted fluorescence microscope captured micrographs using an Andor iXon+ EMCCD camera, an ASI motorized stage, and a shuttered-mercury-lamp light source (X-cite, Lumen Dynamics, Mississauga, Canada).

### Image Analysis

Protein bands in uniform PAG were analyzed as previously described with MATLAB^24^ by creating an array of regions of interest, performing background subtraction, plotting the intensity profile of a band, fitting a Gaussian, performing quality control (signal-to-noise ratio is ≥ 3, *R*^2^ >0.7), and calculating the area-under-the-curve (AUC).^24^ Protein bands in composite gels were analyzed as previously described with MATLAB,^24^ where the median filter (medfilt2 MATLAB function) is applied to reduce noise. Quality controls for all proteins bands are signal-to-noise ratio is >3 and the intensity profile fit a Gaussian curve (*R*^2^ > 0.6).

## Conclusion

We designed SplitBlot as a multimodal single-cell assay that uses arrays of microwells to isolate individual cells, then uses a composite polymer fractionation axis under the action of electrophoresis to perform paired analysis on genomic DNA and protein targets that reside in both the nucleus and the cytoplasm. A key advance is co-measurements of DNA and intracellular nuclear proteins from the same originating single cell. We designed and validated a planar composite agarose-PAG device that, first, functions as a fractionation axis and then toggles into two different molecular analysis workstreams: PCR for nuclear DNA and western blot for nuclear and cytoplasmic proteins. The molecular analysis workstreams employ precise molecular fractionation of DNA and intracellular proteins from single cells via directed electromigration of DNA and proteins along the composite fractionation axis. The device design accommodates molecular fractionation, encapsulation, physical extraction of DNA-laden agarose gel pallets, and pallet transfer for in-tube PCR. The DNA analysis workstream occurs concurrently with the protein analysis workstream which consists of protein fraction-selection, PAGE analysis, and photocapture of protein targets in the PAG region of the axis.

Nuclear proteins, including transcription factors, chromatin proteins, and nuclear structural proteins, play crucial roles in various diseases. This makes multimodal assays capable of measuring nucleo-cytoplasmic proteins valuable for investigating protein regulation mechanisms and identifying disease biomarkers. Furthermore, same-cell single-cell measurements can assess gene editing and examine how gene edits influence nucleo-cytoplasmic protein expression. Current single-cell multimodal assays have a face challenges in nuclear protein analysis due to drawbacks in cell fixation and permeabilization.

The SplitBlot was applied to measure a nuclear protein marker along with genomic (nuclear DNA) by the two concurrent but physically separate analytical work streams (i.e., single-cell western blot and PCR). After running the complete SplitBlot workflow, the assay reported successful amplification of TurboGFP DNA while concomitantly reporting histone H3 expression from the nuclear compartment, from the same originating single cell. Further, cytoplasmic beta-actin protein was also detected by the western blot. Our results confirm that histone H3 protein expression exhibits heterogeneity at the single-cell level and can be detected concurrent with but separately from PCR of the genomic DNA.

## Supporting information

Supplemental dwg file

## Acknowledgement

This work was funded by the Chan Zuckerberg Biohub Investigators Program and by the National Institutes of Health (NIH) R01CA20301 (AEH). Photolithography was performed in the QB3 Biomolecular Nanotechnology Center at UC Berkeley. We acknowledge all lab members of the Herr lab and Paul Lum. U251 cells engineered to express TurboGFP were generously provided by S. Kumar’s Lab. We acknowledge the use of AI-assisted tools were used for language refinement and improving clarity of the text. However, all critical analysis, figure development, and final edits were conducted by authors. All intellectual contributions, interpretations, and conclusions presented in this work remain the responsibility of the authors.

## Supporting Information Available

- Schematics of fabrication of PDMS mold, schematics of fabrication of composite gels, and troubleshooting guide table (PDF)
- CAD files of mask for mold fabrication (.dwg)

## Notes

### Competing Interest Statement

The authors have declared no competing interest.

