## Supplementary Information for "Paired Analyses of Nuclear Protein Targets and Genomic DNA by Single-Cell Western Blot and Single-Cell PCR"

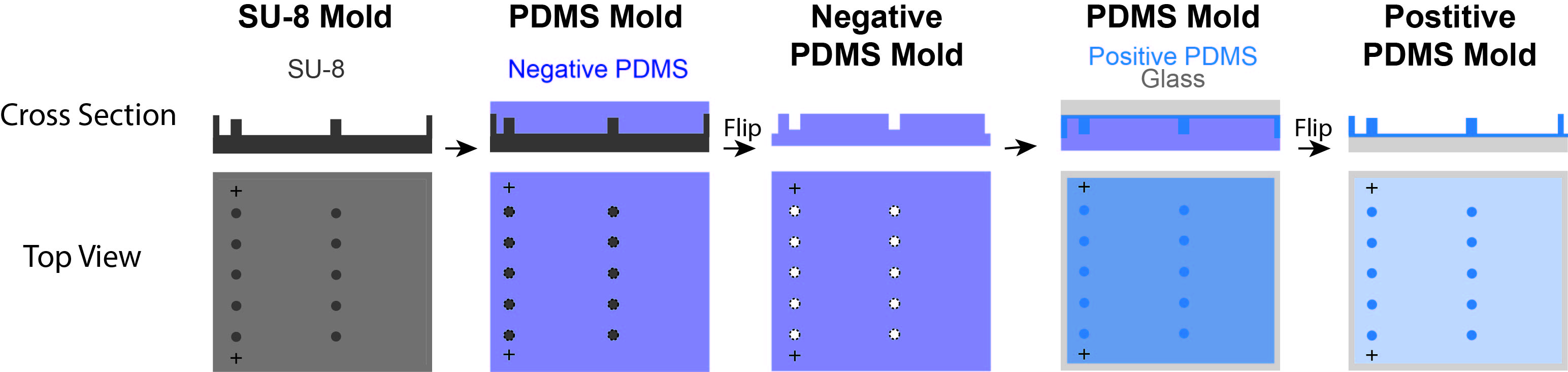

SI Figure 1. PDMS mold fabrication using double mold casting to make 32 μm diameter and 42 μm high microwells. SU-8 was silanized, a negative 10:1 base to curing agent ratio of PDMS was molded and cured at 150 °C for 30 min. The negative PDMS mold was treated with air plasma and silanized for 1hr. The 10:1 PDMS was molded between an air plasma treated glass and the negative PDMS mold. Pressing down on glass removed bubbles. The positive PDMS mold was cured and thermally bonded to the glass at 130 °C for 7 hrs.

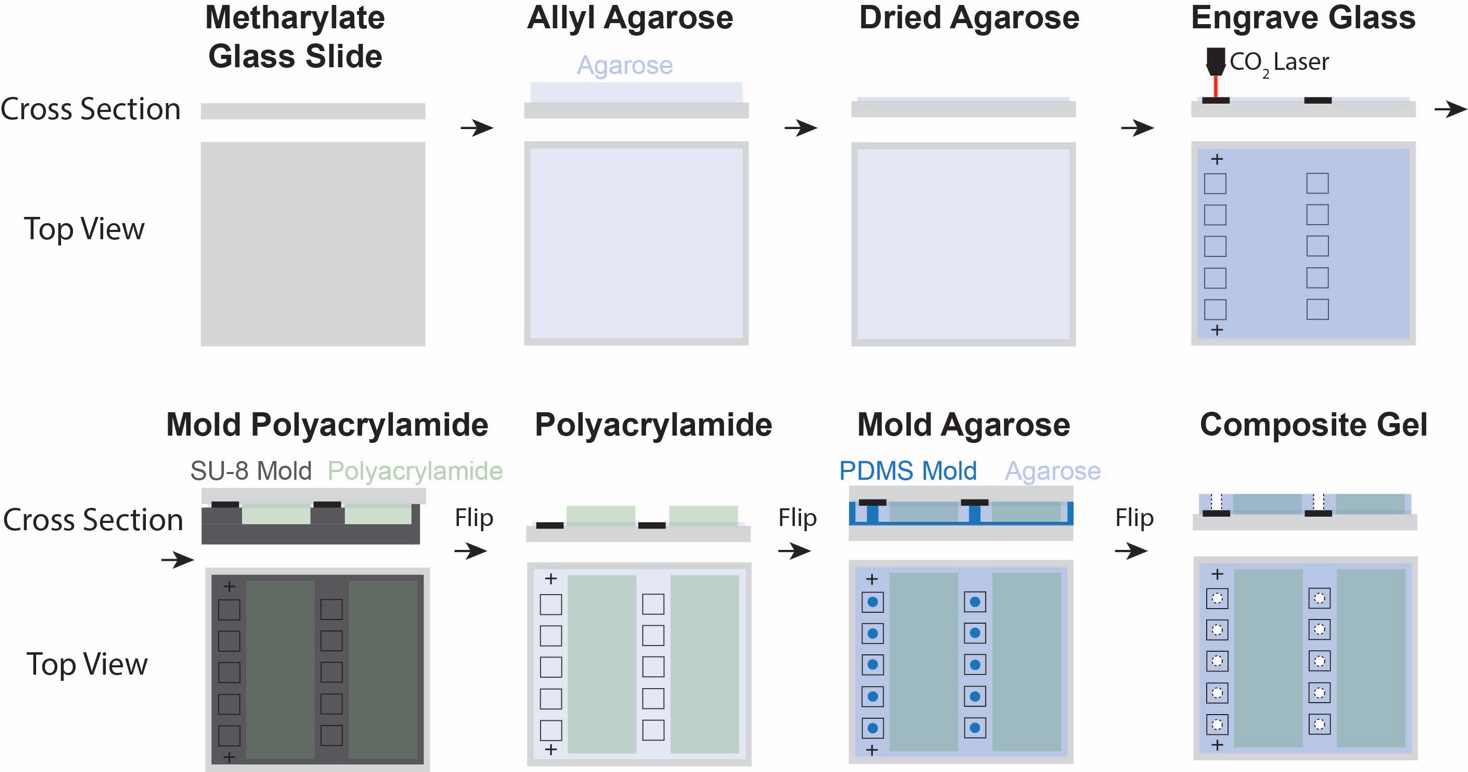

SI Figure 2. Schematic of fabrication of agarose-polyacrylamide composite gel. A methacrylate treated glass slide was coated with a thin layer of molten allyl agarose, the agarose was gelled, and the agarose was air dried. Then, a carbon dioxide laser was used to engrave the pallet region. Polyacrylamide was molded outside the pallet region with a 42 μm high SU8 mold. Agarose microwells 32 μm in diameter and 42 μm high were molded into the pallet region using a PDMS mold.

SI Table 1. Troubleshooting Table

| Observation | Solutions |
| --- | --- |
| No cells in wells | (A) PDMS mold can be damaged resulting in wells that do not isolate cells. Make new PDMS mold and ensure micropost edges are clearly visible under a brightfield microscope. (B) Ensure that agarose remains hydrated before settling and that PBS layer >200μL is on the gel surface before gravity settling cells. |
| No proteins bands in polyacrylamide | (A) Excess Gel Slick in SU-8 mold prevents agarose-polyacrylamide adherence and proteins do not inject at the interface. Dry Gel Slick until it is not visible. (B) Ensure that there is a weight of ~20 g on top of the agarose during molding so wells and polyacrylamide gels are on the same plane. (C) Lysis temp and EP time may be adjusted to detect proteins (SI Figure 3). |
| Proteins bands do not pass quality control in MATLAB analysis | (A) Increase median filter size to reduce noise from the composite gel. (B) Ensure the region of interest dimensions was customized based on the size of protein bands of interest. |
| DNA found outside of microwells | (A) Unsettled cells were lysed outside of microwells. Wash unsettled cells with 15mL of PBS. Repeat until the surface of the gel is clean. (B) Avoid gel dehydration during gravity settling of cells. |

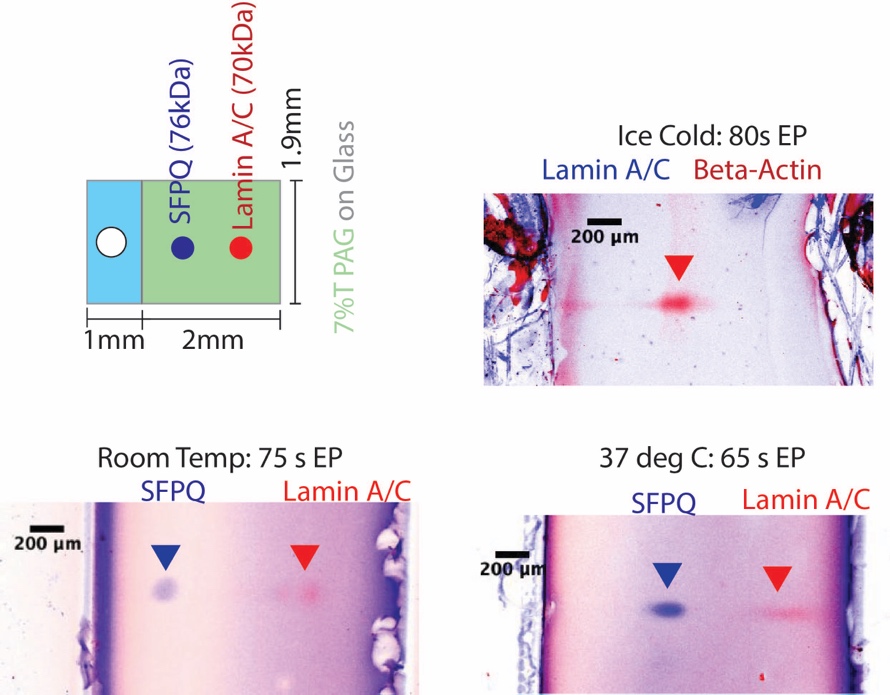

SI Figure 3. After ice cold lysis, beta-actin (42 kDa) was detected but not lamin A/C (70 kDa). Increasing the lysis and EP temperature to room temperature and 37 °C allowed for detection of nuclear proteins Lamin A/C (70 kDa) and SFPQ (76 kDa) in the composite gel. Electric field is 40 V/cm.
